# *Induction of GLUTAMINE DUMPER1* reveals a link between amino acid export, abscisic acid, and immune responses

**DOI:** 10.1101/2020.09.23.310615

**Authors:** Shi Yu, Delasa Aghamirzaie, Kim Harich, Eva Collakova, Ruth Grene, Guillaume Pilot

## Abstract

Amino acid homeostasis in plants is finely tuned to match developmental needs and response to adverse environments. Over-expression of the single-transmembrane domain protein GLUTAMINE DUMPER1 (GDU1) leads to increased amino acid export, reduced growth and constitutive induction of immune responses. We used an inducible gene expression system to tease apart the primary and secondary effects caused by *GDU1*, and demonstrated that the primary effect is increasing amino acid export, followed by increased amino acid content and abscisic acid (ABA) response, and a subsequent activation of defense responses. The *GDU1*-mediated hypersensitivity to ABA partially depended on the E3 ubiquitin ligase LOSS-OF-GDU1 2 (LOG2), a known GDU1 interactor. More importantly, the lysine catabolite pipecolic acid played a pivotal role in the *GDU1*-induced defense responses. This work unravels a novel relationship between amino acid transport, ABA and defense responses, potentially mediated by the GDU1-LOG2 complex, critical for understanding how plants respond to amino acid imbalance.

**ONE SENTENCE SUMMARY:** Genetically induced disturbance of amino acid homeostasis sequentially triggers responses to abiotic stresses and plant defenses to pathogens in Arabidopsis through undefined sensing mechanisms

## INTRODUCTION

Apart from being the building blocks of proteins, amino acids play a central role in plant metabolism along with carbohydrates. Nitrogen enters metabolism through the synthesis of Gln from oxaloacetate, catalyzed by the glutamine synthetase / glutamate synthase cycle (Coruzzi et al., 2015). Amino acids are used for synthesis of specialized metabolites (Pratelli and Pilot, 2014), serve as non-toxic carriers of reduced nitrogen between the organs where they are synthesized to developing tissues. Translocation of amino acids within the plant and across intracellular membranes is mediated by dedicated transporters, which, for the most part, function either as proton-coupled importers or as exporters, and constitute the AAAP, APC and UMAMIT families (Dinkeloo et al., 2018). While the role of several of these transporters is elucidated (Tegeder, 2014), the mechanisms controlling their expression and interactions with metabolic and hormonal pathways remain poorly characterized (Pratelli and Pilot, 2014).

Using a forward genetic screening approach, we identified an unknown protein as being a putative regulator of amino acid export in Arabidopsis. The over-expression of this protein, GLUTAMINE DUMPER 1 (GDU1), led to the development of a pleiotropic phenotype whose most remarkable feature was the secretion of almost pure Gln from the leaves (Pilot et al., 2004). *gdu1-1D* plants are smaller than the wild type, accumulate more amino acids in the leaf, apoplasm, phloem and xylem (Pilot et al., 2004; Pratelli et al., 2010; Pratelli et al., 2012), and display enhanced amino acid export from cells (Pratelli et al., 2010), supporting a role of GDU1 in regulation of amino acid export and homeostasis. Less understood features of *gdu1-1D* plants included constitutive development of necrotic lesions on the leaves and induction of immune responses (Chen et al., 2010; Liu et al., 2010). Suppression of the phenotype, but not protein over-accumulation, by specific amino acid substitutions within the GDU1 sequence (Yu et al., 2015) and by suppression of the activity of LOSS-OF-GDU1 2 (LOG2) (Pratelli et al., 2012) show that this phenotype is not caused by any toxicity effect of GDU1 protein over-accumulation.

GDU1 is a single-transmembrane domain protein with no known functional domain, targeted to the plasma membrane and the endosomal compartments, belonging to a plant-specific family (Pratelli and Pilot, 2006). The conserved cytosolic domain of GDU1 interacts with the membrane-anchored ubiquitin ligases LOG2 and LOG2-LIKE UBIQUITIN LIGASES (LULs) (Pratelli et al., 2012). Both the interaction with LOG2 and the ubiquitin ligase activity of LOG2 are necessary for the development of the Gdu1D phenotype (Pratelli et al., 2012; Guerra et al., 2017), and suppression of *LOG2* expression restores many of the characteristics of the Gdu1D phenotype (Pratelli et al., 2012). In the current model, the GDU1-LOG2 complex is involved in the regulation of amino acid export by targeting an elusive regulator of amino acid exporters for degradation (Guerra et al., 2017). Substrates of LOG2 include GDU1 itself, probably an “incidental” substrate (Guerra et al., 2017), and RESPONSIVE TO DEHYDRATION21 (RD21), a cysteine-type endopeptidase possibly involved in abiotic stress responses (Kim and Kim, 2013). The phenotype of the *LOG2* knockout mutant *atairp3* shows that LOG2 acts as a positive regulator of ABA signaling, but its precise role remains to be defined (Kim and Kim, 2013). The putative connection between ABA and amino acid transport in *log2* is intriguing, because ABA signaling has not been previously linked to amino acid transport.

Cross-talk between phytohormones has been extensively described (Harrison, 2012; Checker et al., 2018), and complex interactions between salicylic acid (SA) and ABA, representing typical biotic and abiotic response pathways, have also been uncovered. The reciprocal effects of ABA on SA are complex, and often appear contradictory and context-dependent: both negative (de Torres Zabala et al., 2009; Manohar et al., 2017) and positive effects (Seo and Park, 2010) have been reported. Little is known about the interactions between ABA and amino acids. It has been shown that ABA and drought can affect amino acid homeostasis both at the mRNA (Less and Galili, 2008; Urano et al., 2009) and amino acid content (Huang and Jander, 2017) levels. Glu or Gln treatment of Arabidopsis leaves and rice roots trigger defense responses using processes partially involving SA (Kadotani et al., 2016; Kan et al., 2017; Goto et al., 2020), and application of a low concentration of Leu induces some defense-related genes in Arabidopsis (Hannah et al., 2010). Despite these studies, the interactions between amino acid metabolism and hormonal signaling pathways are not understood at the molecular level. The amino acid-related and stress-related phenotypes of *gdu1-1D* make the understanding of the role of GDU1 a valuable tool to study this problem.

The characteristics of the Gdu1D phenotype implies metabolic, transport and hormonal alterations. In this study, we sought to establish causality between these effects, notably whether transport alterations were (1) the primary effect of over-expression of *GDU1*, (2) due to disturbances in amino acid homeostasis, or (3) consequences of activation of stress response pathways. In particular, we wanted to assess the role of the ABA pathway and the role of the SA pathway in the Gdu1D phenotype. To answer these questions, we recapitulated the Gdu1D phenotype caused by over-expression of *GDU1* using an inducible gene expression system. After induction, the development of the phenotype was closely followed over time at the molecular, metabolic and physiological levels. The results allowed us to differentiate primary and secondary effects caused by overexpression of *GDU1* and infer causality between various phenotypes.

## RESULTS

### Induction of GDU1 recapitulates the Gdu1D phenotype

Because the *gdu1D* mutation caused a gain-of-function phenotype (i.e., constitutive overexpression), we reasoned that inducible overexpression of the wild-type gene could recapitulate phenotypes of the *gdu1-1D* mutant. Sampling over time, following induction of *GDU1*, would enable us to determine the temporal sequence of events following induction of *GDU1*. Three independent Arabidopsis lines that express the *GDU1* gene under the control a dexamethasone-inducible promoter (pOp/LhGR system (Craft et al., 2005; Samalova et al., 2005), were constructed and brought to homozygosity (lines DEX1, DEX2, DEX3; Supplemental Text 3). Mature plants were sprayed with dexamethasone and studied over time. *GDU1* mRNA accumulation peaked between 6 and 12 h post induction (HPI), accumulating ∼5,000 to ∼10,000 times more than in the 4c-S7 control line (Figure 1A), an amount higher than in the constitutive *gdu1-1D* and *gdu1-2D* mutants (∼500 and 250 times over-accumulation, respectively; Yu and Pilot, unpublished data). *GDU1* mRNA leveled off after 24 h to approximately 3,000 times the level of 4c-S7 until the end of the experiment.

**Figure 1.**
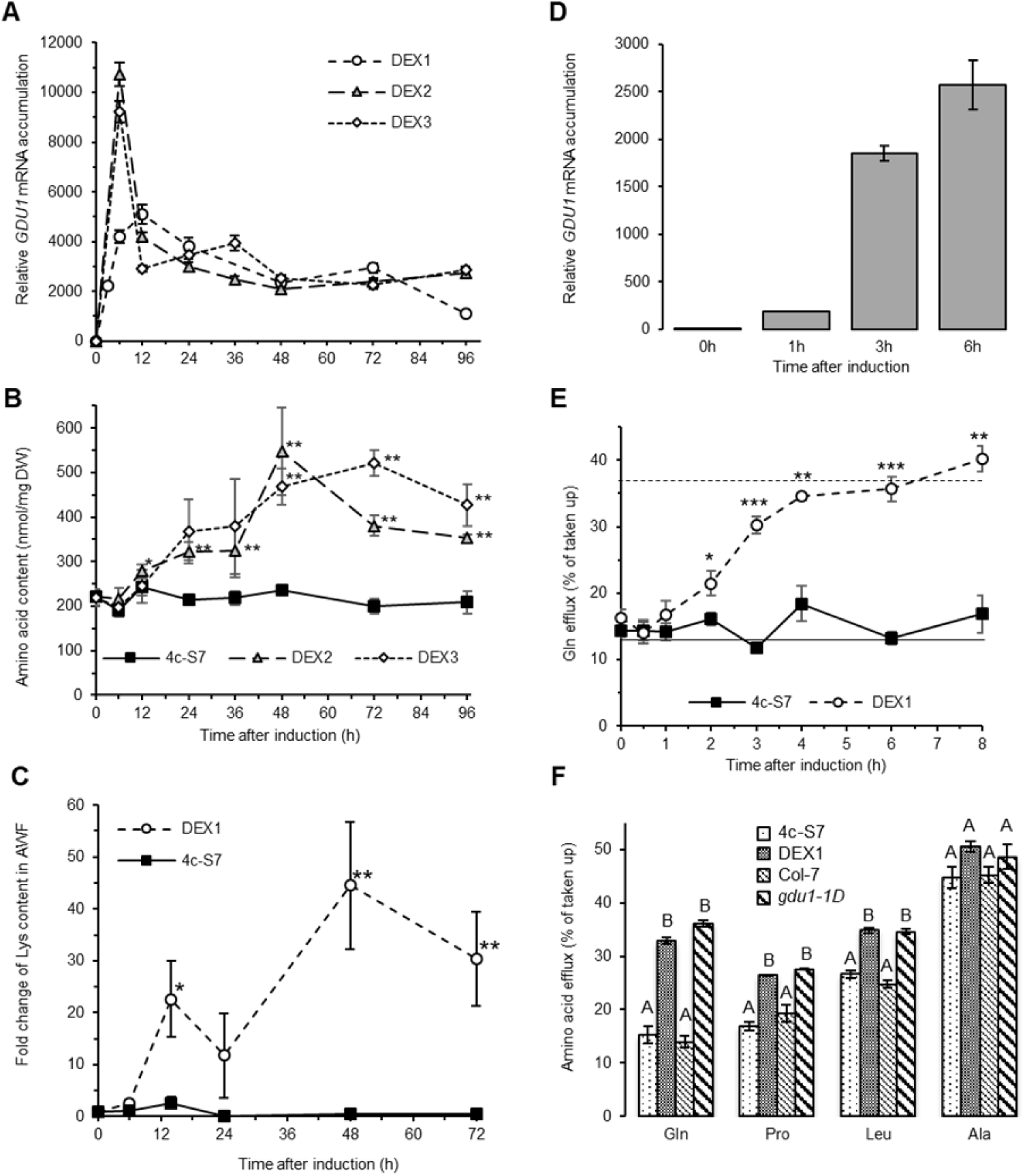
Time course analysis of *GDU1* mRNA accumulation, amino acid content, and amino acid export after induction of *GDU1*. **A**. Lines DEX1, DEX2, DEX3 and the pBIN‐LhGR‐N activator line 4c‐S7 (control line) were grown for three weeks on soil, and sprayed with dexamethasone. *GDU1* mRNA accumulation was measured by qPCR over the course of four days after induction. Line DEX1 was tested in a separate experiment as lines DEX2 and DEX3. Relative accumulations are reported as fold change compared to the 4c-S7 line at each time point, set at 1. Error bars=SE, n=3 biological replicates. **B**. Free amino acid content in whole leaves from the same samples as in A. Error bars=SE, n=3 biological replicates. Statistically different from the 4c-S7 line: t-test (* p<0.1; ** p<0.05). **C**. Change in Lys content in apoplasmic wash fluid (AWF) of lines DEX1 and 4c-S7 grown for four weeks on soil and sprayed with dexamethasone. Content was reported to the content at time 0 for each line, set at 1. Error bars=SE, n=4 biological replicates. Statistically different from the 4c-S7 line: t-test (* p<0.1; ** p<0.05). **D, E and F**. The 4c-S7 and DEX1 lines were grown on half strength MS +1% sucrose for one week, followed by four additional days in liquid medium, and treated with 30 µM dexamethasone for times indicated on the graphs. **D**. Fold change in *GDU1* mRNA content in the DEX1 line compared to the 4c-S7 line at each time point (set at 1). Error bars = SE, n=3 biological replicates. **E**. Measurement of Gln efflux over time after *GDU1* induction. Statistically different from the 4c-S7 line: t-test (* p<0.05, ** p<0.01, *** p<0.001). The dashed and plain horizontal lines correspond to the Gln efflux of *gdu1-1D* and Col-7 respectively, measured in the same conditions. Error bars=SE, n=3-6 biological replicates. **F**. Efflux Gln, Pro, Leu, and Ala (measured for 20 min after 20 min uptake of 1 mmol.l^-1^ of each amino acid) at three hours after dexamethasone treatment on the 4c-S7 and DEX1 lines, and in Col-7 and *gdu1-1D* lines. Different letters indicate significantly different results for each amino acid (ANOVA-Tukey, p<0.05). Error bars=SE, n=3 biological replicates.

To determine to which extent inducing *GDU1* expression recapitulates the Gdu1D phenotype, free amino acid content of leaves of the DEX2 and DEX3 lines was measured from the same plants as above. Amino acid levels started to increase after 12 HPI, being 50% higher at 24 h and about 100% higher from 36 HPI compared to 4c-S7 (Figure 1B, Supplemental Table 1). Most amino acids, except for Ala, Asp and Glu, were responsible for this increase (Supplemental Table 1). Other than Lys (15 fold increase) the amino acid levels increased 2-4 fold compared to the control at 96 HPI, and largely mirrored the amino acid levels of constitutive *GDU1* over-expressors examined in a previous study (Pilot et al., 2004). Total free amino acid levels in the apoplasm wash fluid were similar for the DEX1 and 4c-S7 plants (Supplemental Table 2), but levels of many amino acids increased at 6 HPI, and stayed elevated until at least 48 HPI (Asn, Gln, Gly, His, Ile, Leu, Phe, Pro, Ser, Thr and Val) or decreased (Asp, Glu and GABA) (Supplemental Table 2). The increase of Lys concentration was dramatic, about 75-fold compared to 4c-S7 at 48 HPI (Figure 1C; Supplemental Table 2). To test if the increase in apoplasmic amino acid concentration resulted from modification in amino acid transport, DEX1 plants were grown in liquid culture, treated with dexamethasone, and assayed for amino acid export. In these growth conditions, dexamethasone induced *GDU1* mRNA accumulation with a similar intensity and kinetics as for soil-grown plants (Figure 1D). Gln efflux of DEX1 started to increase significantly at 2 HPI, reaching levels similar to the constitutive over-expressor *gdu1-1D* (30% vs. 36%) by 3 HPI (Figure 1E). Induction stimulated Pro and Leu export to the same extent as in *gdu1-1D* (Figure 1F).

White crystalline deposits were observed as soon as 3 days after dexamethasone induction (Figure 2B), and their size increased over time (data not shown), mimicking the Gln deposits at the hydathodes of the *GDU1* over-expressors (Pilot et al., 2004). To test whether dexamethasone-induced plants would display tolerance to toxic concentrations of amino acids similarly to the *GDU* over-expressors (Pratelli and Pilot, 2007; Pratelli et al., 2010), plants were induced and grown on sterile medium containing 4 mM Ile. Roots of the DEX lines grew as well as *gdu1-1D* in presence of Ile and dexamethasone, while growth was inhibited similarly to Col-0 by Ile in absence of dexamethasone (Supplemental Figure 1).

**Figure 2.**
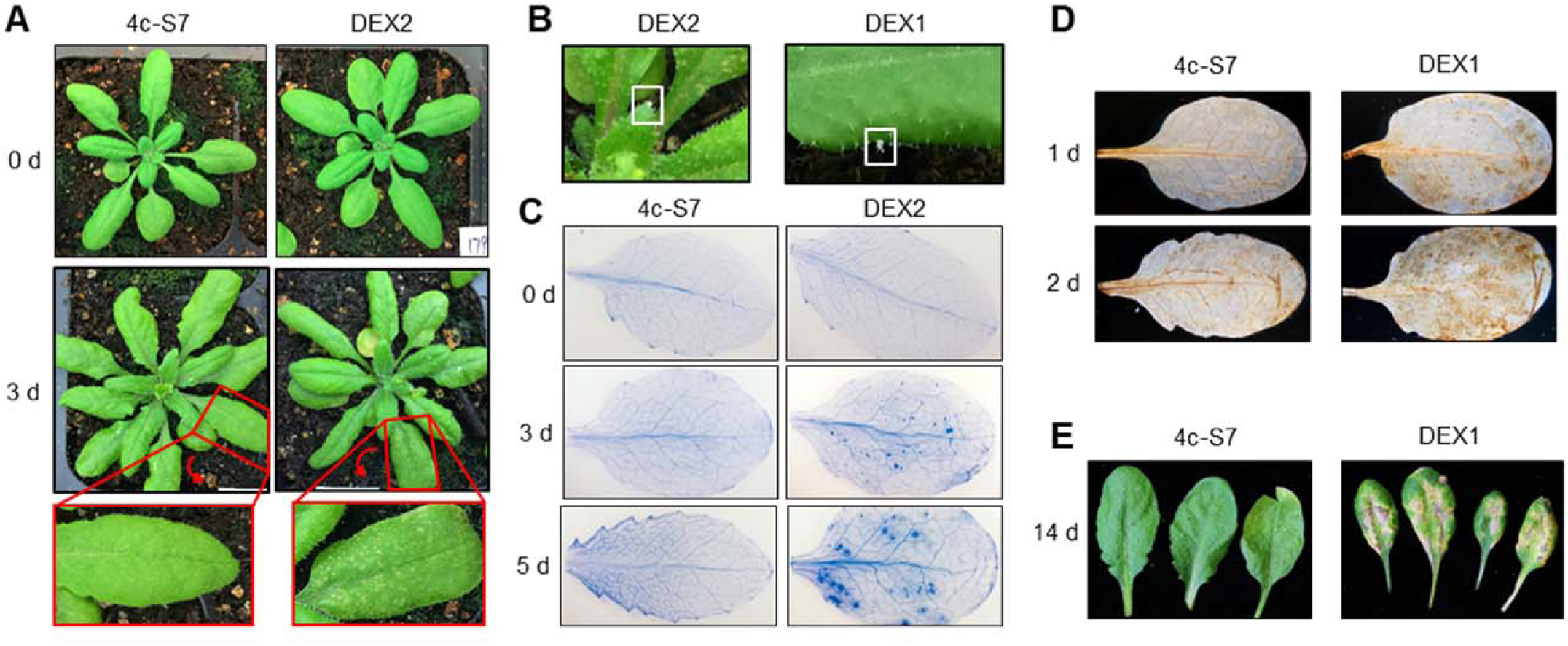
Phenotype of the DEX and 4c-S7 plants after spraying with dexamethasone. **A**. Picture of DEX2 and 4c-S7plants at 0 and 3 days after induction. Red boxes: enlargement of the leaves at 3 days, showing the development of lesions in DEX2 only (arrows indicate the direction of rotation). **B**. Close-up view of leaves of DEX1 and DEX2, three days after induction, showing Gln secretion (white boxes). **C**. Leaves of 4c-S7 and DEX2 plants, 0, 3 and 5 days after induction, stained using Trypan blue to reveal cell death. **D**. Leaves of 4c-S7 and DEX1 plants, 1 and 2 days after induction, stained with diaminobenzidine to reveal presence of reactive oxygen species. **E**. Leaves of 4c-S7 and DEX1 plants, 14 days after induction (induction was repeated 7 days after the first spray).

Similar to the constitutive over-expressor *gdu1-1D* (Supplemental Figures 2A,B), necrosis spots on the whole area of the leaves were observed after three days post-dexamethasone induction on all DEX lines, but not on the 4c-S7 line (Figure 2A, insert). Such lesions can be a sign of cell death, which is often associated with the presence of reactive oxygen species (ROS) (Van Breusegem and Dat, 2006). Accordingly, ROS and cell death were detected after staining DEX1 leaves with 3,3’-diaminobenzidine (DAB) and Trypan blue, respectively (Pogany et al., 2009): ROS accumulated as soon as 24 HPI (Figure 2D), while necrotic spots appeared after 3 days post induction (Figure 2C), and were comparable both in size and number to *gdu1-1D* (Supplemental Figure 2C). When DEX1 plants were sprayed twice with dexamethasone seven days apart to prolong the induction for 14 days, leaves of the DEX1 plants displayed extensive yellowing and parched areas, very similar to the older leaves of *gdu1-1D* (Figure 2E and Supplemental Figure 2A).

Altogether, these results demonstrate that induction of *GDU1* using the inducible pOp/LhGR system leads to a rapid and robust expression of *GDU1*, and recapitulates in about 3-4 days the most identifiable characteristics of the Gdu1D phenotype, namely increased amino acid export, tolerance and content, as well as development of lesions.

### Induction of GDU1 activates ABA and defense pathways in a sequential way

Lesions are often associated with cell death mediated by SA (Nimchuk et al., 2003), while long term ABA treatment is known to induce leaf yellowing (Wang et al., 2018). To determine the effect of *GDU1* induction on the activity of the SA and ABA pathways, mRNA accumulations of biosynthesis (*SID2, NCED3*) and response (*PR1, RD29A*) marker genes for SA and ABA respectively were measured over time. The kinetics of induction for each gene was slightly different between the DEX lines but followed the *GDU1* induction kinetics, and the same trends were observed: *NCED3* mRNA peaked first at about 12 HPI, followed by *RD29A* (12-24 HPI), *SID2* (24-48 HPI) and *PR1* (48-72 HPI) mRNAs (Figures 3A-D; Supplemental Figure 3A and 3B). The slower response of the DEX1 line upon induction allowed us use other marker genes to elucidate the induction of SA (*ICS2, PAL4*), ABA (*KIN1, COR15, CBF3, RAB18*), auxin (*IAA5*), jasmonate (*OPR3, PDF1*.*2A*) and ethylene (*ERS2*) pathways (Supplemental Table 3 for details of the genes). The response to ABA and SA was confirmed (Supplemental Figures 3A and 3B), while no strong and durable ethylene or auxin responses were detected. Synthesis of and response to jasmonate were induced from 24 HPI and 72 HPI respectively (Supplemental Figure 3C). In good agreement with the results of the marker gene study, ABA and JA contents peaked at 24 HPI and declined back to the levels of 4c-S7 after 48-72 HPI (Figures 3E and 3F), while SA steadily accumulated over the time of the induction from 48 HPI (Figure 3G).

**Figure 3.**
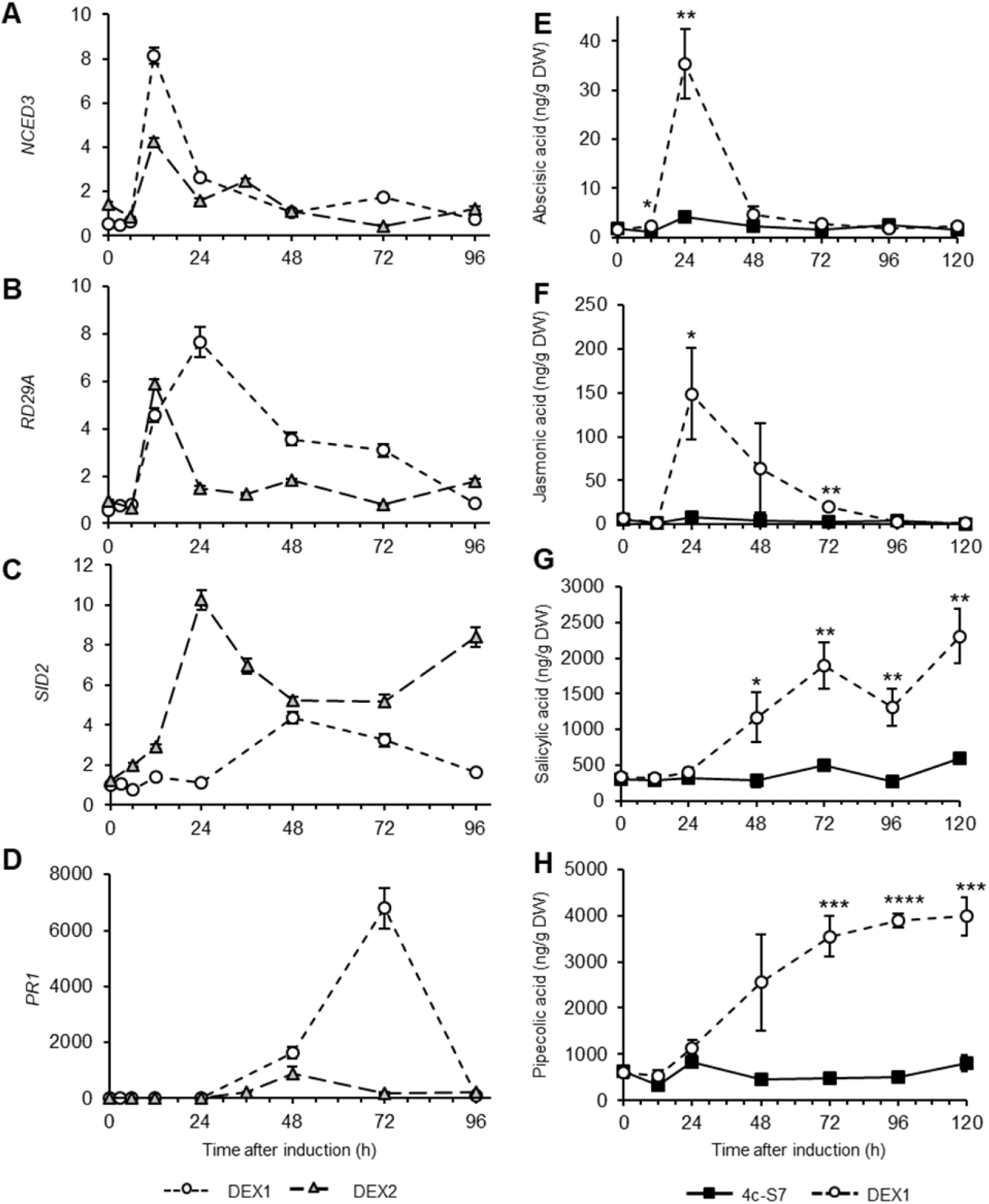
Time course analysis of the expression of marker genes and of the accumulation of hormones following *GDU1* induction. **A to D**. DEX1 and DEX2 lines were induced in two separate experiments reported in Figure 1A. Data represent fold difference of the mRNA accumulations of each gene, relative to the corresponding mRNA content in the control 4c-S7 line at the corresponding time point, set at 1. *NCED3* (**A**) and *RD29A* (**B**), markers for ABA synthesis and signaling; *SID2* (**C**) and *PR1* (**D**), markers for SA synthesis and signaling, respectively. Raw data are presented in Supplemental Figure 3. Error bars are SE, N=3 biological replicates. **E** to **F**. 4c-S7 and DEX1 plants were grown for three weeks on soil, sprayed with dexamethasone. Hormone content was measured by LC-MS. (**E**) Abscisic acid; (**F**) Jasmonic acid; (**G**) Salicylic acid; (**H**) Pipecolic acid. Error bars=SE, n=4 biological replicates. Statistically different from 4c-S7: t-test (* p<0.05, ** p<0.01, *** p<0.001, **** p<0.0001).

Pipecolic acid (Pip) is a transported compound synthesized from Lys, necessary for the establishment of systemic acquired resistance (SAR) (Navarova et al., 2012; Bernsdorff et al., 2016). Since Lys accumulates to high levels upon *GDU1* induction, the content of the mRNAs corresponding to the enzymes that catalyze the degradation of Lys and the conversion of Lys to Pip (*LKR-SDH* and *ALD1* genes respectively) were measured in the DEX1 line. *LKR-SDH* mRNA increased at 24 HPI, while *ALD1* mRNA accumulation peaked at 48 HPI with a remarkable 350-fold increase (Supplemental Figure 3D). At the same time, Pip content steadily increased over time, mirroring SA accumulation (Figure 3H). Interestingly, these hormonal responses occurred sequentially, well after the onset of the increase in amino acid export (2 HPI). Measurement of Gln uptake from plants treated by ABA, SA or Pip further proved that the increase in amino acid export was not triggered by these phytohormones (Supplemental Figure 4).

**Figure 4.**
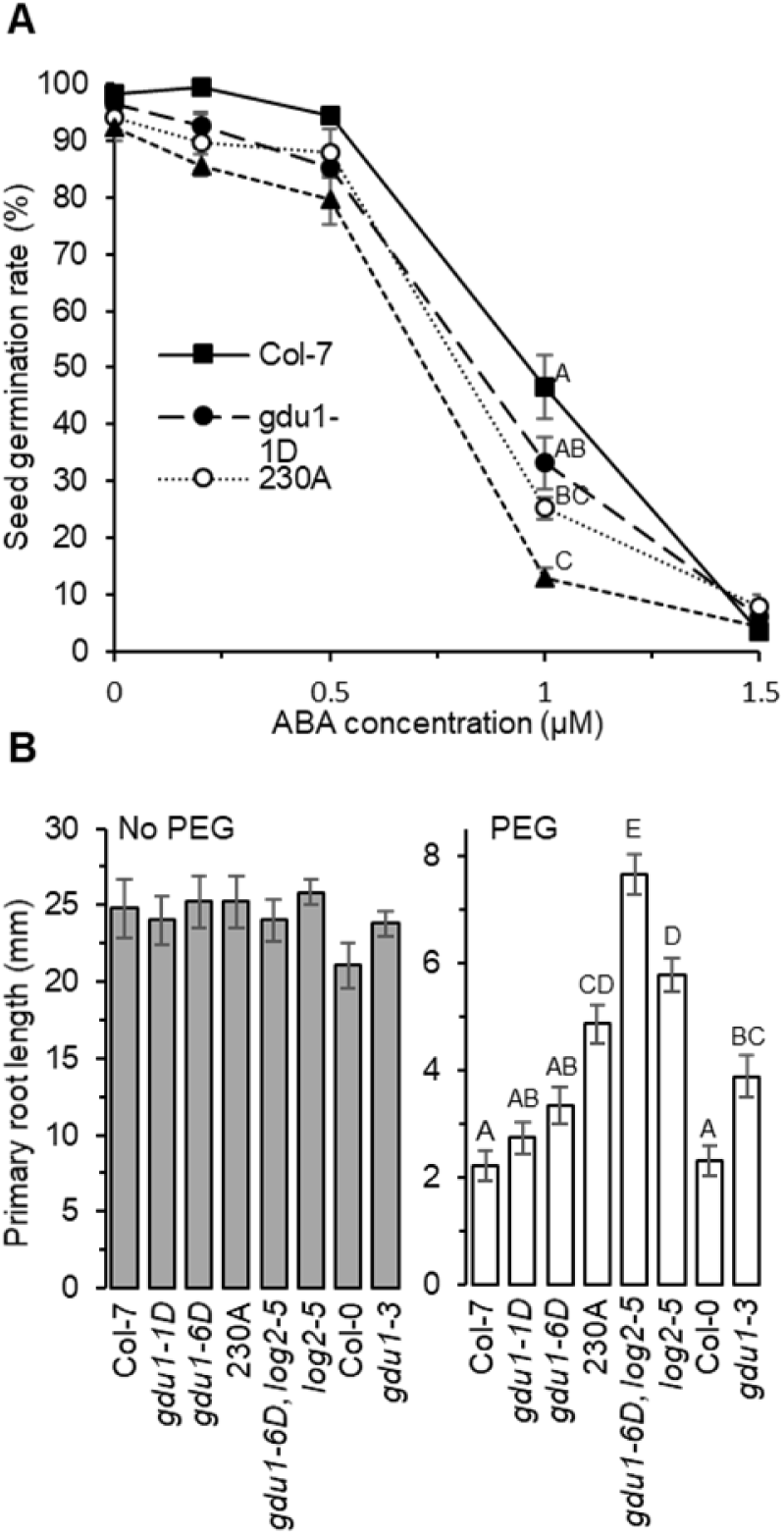
Sensitivity to ABA and drought of plants altered in *GDU1 and/or LOG2* expression. **A**. Germination rate of Col-7, *gdu1-1D, gdu1-6D* and *gdu1-1D* expressing an amiRNA targeting *LOG2* (230A). Plants were grown for five days on half strength MS with 1% sucrose containing various concentrations of ABA indicated in the figure. Germination was assessed as cotyledon greening. Error bars=SE, n=3 biological replicates consisting of 60-240 seedlings each. **B**. Length of the primary root of plants grown for seven days on PEG-containing (mimic drought) or PEG-free (normal) medium. *gdu1-3* is a T-DNA knockout mutant of GDU1, *log2-5* is a suppressor mutation of the Gdu1D phenotype of line *gdu1-6D* (Pratelli and Pilot, unpublished). Error bars=SE, n=18. Different letters for the PEG conditions indicate significantly different results (ANOVA-Tukey HSD, p<0.05).

An unbiased transcriptomics approach confirmed our qPCR results, in that the primary responses of the transcriptome to *GDU1* induction is stress signaling, followed by ABA responses, and defense responses (Supplemental Text 1). Mining the genes induced at 7 h after induction did not uncover a specific pathway which could help explain the phenotype related to amino acid transport (Supplemental Files 2 and 3). We mainly found genes associated with the general GO terms “Regulation of transport”, and “Response to stimulus”, suggesting that, at that time point, the plant is initializing responses whose nature cannot be deduced from the data.

### GDU1-induced hypersensitivity to ABA partially depends on LOG2

Since the ABA signaling pathway is induced by *GDU1* over-expression, we hypothesized that *GDU1* over-expressor, but not the *gdu1-3* knockout mutant, should therefore exhibit hypersensitivity to exogenous ABA. We observed that *GDU1* overexpressors (Pratelli and Pilot, 2006), but not *gdu1-3*, were indeed hypersensitive to ABA in a germination assay (Figure 4A; Supplemental Figure 5). The LOG2 ubiquitin ligase is a positive regulator of ABA signaling (Kim and Kim, 2013) and its activity is stimulated upon interaction with GDU1 (Pratelli et al., 2012; Guerra et al., 2017). To test the involvement of LOG2 into the GDU1-mediated ABA response, *LOG2* expression was reduced by expressing an artificial miRNA targeting *LOG2* in the *gdu1-1D* background (line 230A (Yu and Pilot, 2014)). The germination rate in presence of ABA of this line was not different from the mutant rate (Figure 4A), suggesting that LOG2 is not indispensable in this specific ABA assay. On the contrary, lines in which *LOG2* or *GDU1* expression was suppressed or reduced, were less sensitive to simulated drought, which is an ABA-dependent process (Rowe et al., 2016), than wild type roots, while the *GDU1* over-expressors *gdu1-1D* and *gdu1-6D* displayed similar root length as the wild type (Figure 4B). Interestingly, root growth of lines 230A and *gdu1-6D log2-5* was less inhibited by drought than both the wild type and *gdu1-1D*, suggesting that knockdown/knockout of LOG2 activity is epistatic to *gdu1-1D* over-expression in this assay, but not in the ABA germination assay. The different results from these assays hints at GDU1 over-expression triggering LOG2-dependent and LOG2-independent ABA responses. In addition, LOG2 is likely not the only gene involved in the GDU1-mediated changes in amino acid transport and homeostasis: while suppression of LOG2 activity in *GDU1* over-expressors brought free amino acid accumulation to wild type levels (Supplemental Figure 6), Gln uptake and efflux were still altered (Supplemental Figure 7). At the same time, suppression of *GDU1* or *LOG2* expression alone, did not affect free amino acid content (Supplemental Figure 6), which indicate that other GDUs (Pratelli and Pilot, 2006) or genes similar to LOG2 (Pratelli et al., 2012) may also be involved in the process and functionally complement those loss-of-function mutants. Therefore, the *GDU1*-induced effects on amino acid transport and content are not entirely dependent on LOG2, similar to *GDU1*-induced ABA responses reported above.

**Figure 5.**
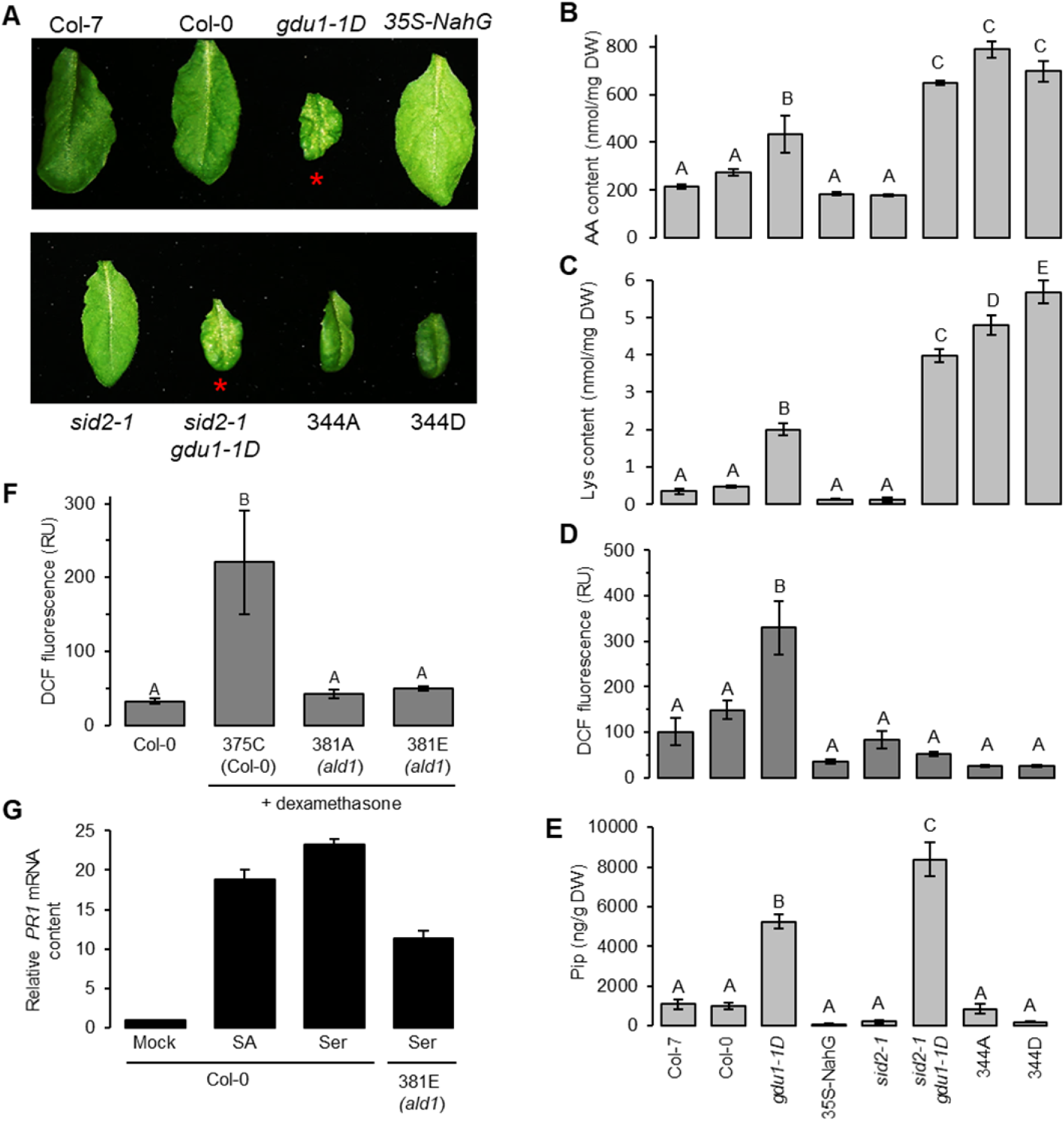
Modification of the phenotype of *gdu1-1D* plants by genetic suppression of defense responses. Wild type (Col-0, Col-7), *gdu1-1D, sid2-1* and the corresponding double mutant, plants over-expressing *NahG* (35S-NahG), and two independent lines coming from the transformation of *gdu1-1D* with the CsVMV-NahG construct (344A and 344D), were grown for five weeks on soil. **A**. Picture of a typical leaf from each plant. Red asterisks indicate leaves with lesions. **B**. Total free amino acid (AA) content in leaves of the plants. Error bars=SE, n=4, see also Supplemental Table 3. **C**. Free Lys content in leaves of the plants. Error bars=SE, n=4. **D**. Reactive oxygen species levels in leaves of each of the lines; RU, relative units. Error bars=SE, n=3-8. **E**. Pip content in leaves of each of the lines. Error bars=SE, n=4. Different letters indicate significantly different results (ANOVA-Tukey HSD, p<0.05). **F**. Col-0 and *ald1* were transformed with a construct leading to the dexamethasone-inducible expression of *GDU1* (see Methods), to yield line 375C (Col-0), and two independent lines 381A and 381E (*ald1*). Lines were grown for four weeks on soil, sprayed with dexamethasone (+DEX) and leaf samples were collected after five days for ROS quantitation. Error bars=SE, n=4-12, different letters indicate significantly different results (Kruskal–Wallis - Dunn’s tests, p<0.1). **G**. Plants were grown for eight days in half strength MS + 1 % sucrose and three additional days in liquid medium. Ser was added to the medium to a final concentration of 5 mM and samples were taken after 2 days. Treatment with 5 µM SA for 2 days was used as a positive control. Fold change in *PR1* mRNA content is expressed compared to the accumulation in Col-0 plants treated with 0.025% DMSO, set at 1. Error bars=SE, n=4 biological replicates.

**Figure 6.**
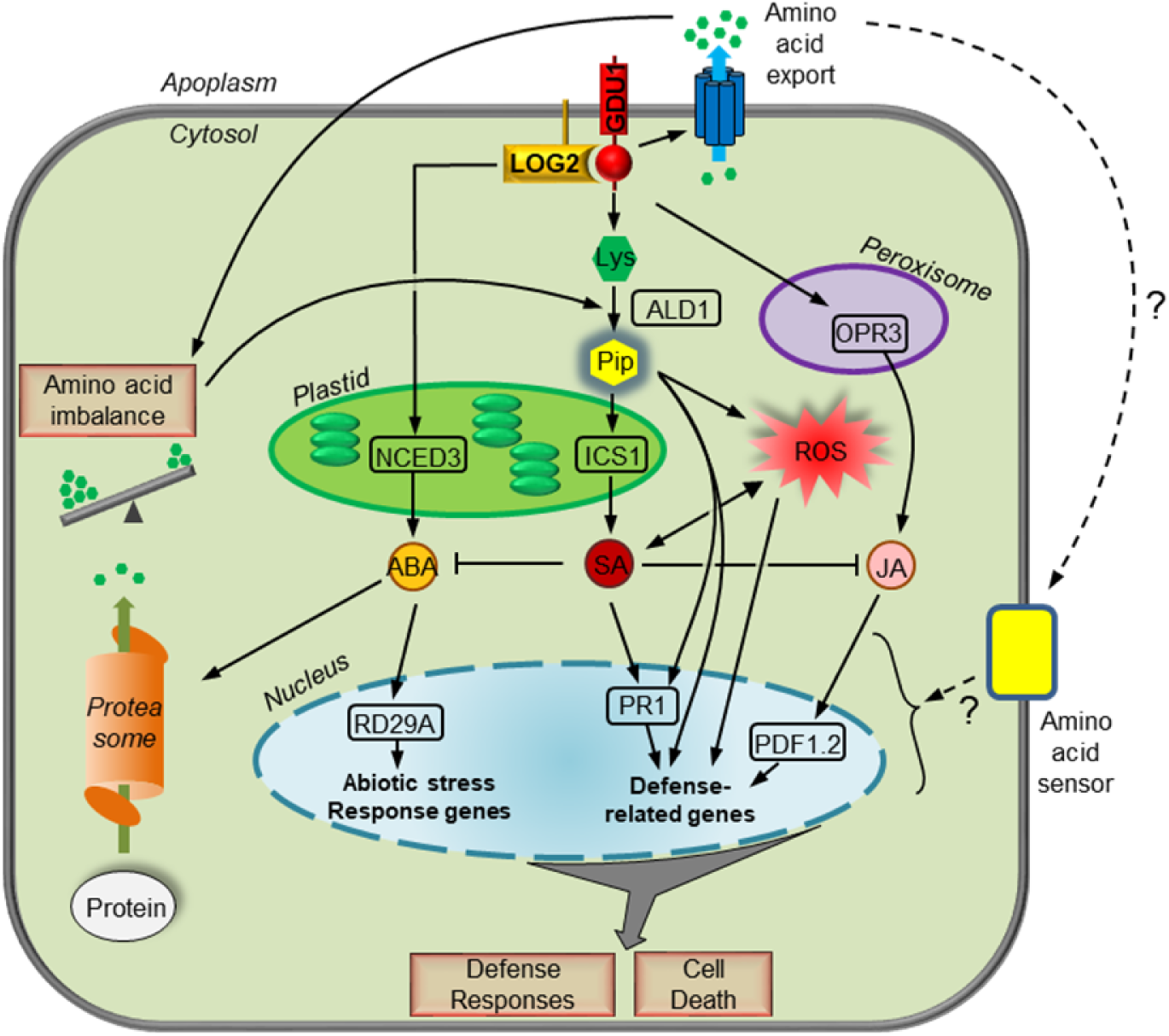
Model for the processes activated upon *GDU1* over-expression. Boxed genes correspond to the main marker genes assayed for expression in Figure 3 and Supplemental Figure 3. Arrows represent activation, and bars represent inhibition. Solid lines indicate confirmed links from this and previous studies, and dashed lines indicate possible links. See main text for details.

### Pip plays a pivotal role in the GDU1-induced defense responses

In addition to developing lesions on leaves, *gdu1-1D* is about 50% smaller than wild type plants (Pilot et al., 2004), a phenotype which could result from constitutive activation of the SA pathway, similar to the *cpr* and *cep* mutants (Bowling et al., 1994; Silva et al., 1999; Gou et al., 2009; Mosher et al., 2010). To test for any causality between SA responses, smaller size and lesion development, SA levels were genetically decreased by crossing *gdu1-1D* plants with the *sid2-1* mutant, in which SA biosynthesis is dramatically reduced (Wildermuth et al., 2001), or by expressing the SA-degrading enzyme *NahG* (Delaney et al., 1994) (lines 344A and 344D). Presence of the *sid2-1* mutation or the NahG protein did not affect the expression of *GDU1* (Supplemental Figure 8), ABA levels (Supplemental Figure 9A) or Gln export (Supplemental Figure 10), but expectedly decreased the *PR1* mRNA accumulation (Supplemental Figure 8) and the content of SA and JA (Supplemental Figures 9B and 9C). However, the *sid2-1* mutation or expression of *NahG* did not restore the size defect of *gdu1-1D* (Figure 5A; Supplemental Figure 11A). Similarly, levels of some amino acid, especially Lys, remained elevated in leaves of those lines (Figures 5B, 5C; Supplemental Table 3) and Gln was still secreted to *gdu1-1D* levels (Supplemental Figure 10). The activation of the SA pathway does not seem to be the cause of the reduced size and the increase in amino acid accumulation and secretion.

Interestingly, while the levels of ROS were reduced to identical or less than wild type levels by expression of *NahG* or the presence of the *sid2-1* mutation (Figure 5D), only expression of *NahG* suppressed the development of spontaneous cell death and lesions (Figure 5A, Supplemental Figure 11B). NahG activity has been shown to suppress the activation of both SA-dependent and SA-independent defense pathways (Heck et al., 2003), implying that over-expression of *GDU1* activates both pathways, the latter being critical for the development of lesions in *gdu1-1D*.

Intrigued by this result, we sought to assess the involvement of Pip, whose content is elevated by *GDU1* over-expression. Compared to the wild type, Pip accumulation was elevated in the *gdu1-1D* and *sid2-1 gdu1-1D* mutants (even to a higher level than in *gdu1-1D*), while it was reduced to wild type levels by expression of *NahG* (Figure 5E). Accumulation of Pip effectively correlated with the presence of lesions (Figure 5A), but the content of Lys, precursor of Pip, did not, as it was enhanced by the *sid2-1* mutation or NahG enzyme (Figure 5C). To confirm the role of Pip in the Gdu1D phenotype, the *ald1* mutant, showing no α-aminotransferase activity (Navarova et al., 2012), and wild type plants were transformed with a construct enabling the induction of *GDU1* by dexamethasone. Compared to the induction of *GDU1* expression in the wild type, inducing *GDU1* expression in the *ald1* mutants did not increase ROS accumulation (Figure 5F), and led to a lower *PR1* mRNA accumulation than in wild type plants (Supplemental Figure 12A), even if *GDU1* expression remained elevated (Supplemental Figure 12B). These results suggest that Pip is necessary for ROS accumulation, lesion development and the induction of SA-dependent and SA-independent pathways in the Gdu1D phenotype.

### Amino acid treatment triggers plant defense responses in a Pip dependent manner

Seeking a possible link between the GDU1-triggered increase in amino acid content and the subsequent activation of plant defense responses, plants expressing GUS under the control of the *PR1* promoter were grown in liquid medium and GUS activity was detected after treatment with SA, Pip, amino acids, or amino acid derivatives. Plants treated with 500 µM Cys, Gln, Gly, Phe, Pro, Ser and Tyr showed a marked increase in PR1-GUS activity (Supplemental Figure 13A) in a dose-dependent manner, similar to Pip (Supplemental Figure 13B-D). This result was confirmed by a 20-fold increase in accumulation of *PR1* mRNA in response to Ser in wild type plants, which was reduced by 50% in *ald1* compared to the wild type (Figure 6G), showing that Pip contributes to the responses to amino acids.

GLUTAMATE RECEPTOR LIKE (GLR) proteins have been proposed to act as amino acid sensor candidates (Forde and Roberts, 2014; Gent and Forde, 2017). We tested the hypothesis that GLRs sense the amino acid secretion triggered by *GDU1* induction and trigger stress responses, as shown upon wounding (Toyota et al., 2018) (Supplemental Text 2). Our results establish that GLRs are not positive regulators of the signaling linking amino acid imbalance and defense responses upon *GDU1* over-expression, leaving Pip and SA as the most important players.

## DISCUSSION

Based on the phenotype of the corresponding over-expressors, the GDU1 protein was previously proposed to be a direct or indirect regulator of amino acid export from plant cells (Pilot et al., 2004; Pratelli et al., 2010). In the current model, GDU1 and the ubiquitin ligase LOG2 control amino acid transport through ubiquitination of (an) unknown substrate(s) (Guerra et al., 2017). Activation of LOG2 upon interaction with GDU1 would lead to a rapid degradation of an inhibitor of amino acid exporter(s), causing an immediate increase in amino acid export. Other possible targets of the GDU1-LOG2 complex could be proteins involved in amino acid/nitrogen sensing and signaling, which, upon interaction with the complex, would trigger post-translational events leading to rapid changes in amino acid transport at the plasma membrane. The present work, initially aimed at finding out how the characteristics of the Gdu1D phenotype are linked to modification of amino acid transport, unraveled intriguing relationships between amino acid transport, metabolism, stress responses and immunity.

### The primary effect of GDU1 over-expression is an increase in amino acid export, caused by post-translational processes

It could be postulated that protein over-accumulation, leading to aggregation and cell death (Ueno et al., 2019), could be the reason of the observed stress and immune responses in *gdu1-1D* and the DEX lines. This hypothesis is invalidated by the observation that specific amino acid substitutions within the GDU1 sequence (Yu et al., 2015) or suppression of the activity of LOG2 (Pratelli et al., 2012) suppress the stress responses without affecting protein accumulation. Studying the effect of the induction of *GDU1* expression over time unequivocally showed that the first event is an increase in amino acid export by 2 HPI (Figures 1E and 1F), with no change in any other parameter detected at that early time point. To trigger the Gdu1D phenotype, *GDU1* mRNA needs to accumulate to levels over 100 folds that of wild type plants (Pilot et al., 2004), an amount reached at about 1 HPI (Figure 1D). Based on an average synthesis rate of 5 amino acids per second and a reasonable trafficking time from the ER to the plasma membrane of 60 min (Hirschberg et al., 1998), the 158 amino acid-long GDU1 protein could be synthesized, folded, and accumulate at the plasma membrane in a little over an hour after being transcribed. Taking into account a protein synthesis time of 30 sec and a half-life time of an hour (Guerra et al., 2017), one can calculate that the accumulation of GDU1 protein at the plasma membrane necessary for triggering the Gdu1D phenotype is reached in about 3 hours after *GDU1* induction (data not shown). This value is in good agreement with the experimental data, which show that amino acid export reaches the level of the constitutive over-expressor at about 3 HPI (Figures 1E and 1F). The phenomenon triggered by *GDU1* induction is thus temporally concomitant with the accumulation of GDU1 at the plasma membrane, supporting the current model that the primary role of GDU1 is to control the activity of the LOG2 ubiquitin ligase (Guerra et al., 2017). Importantly, no *de novo* transcription or translation of other genes following induction would be necessary to explain this effect. This would explain why no signature or pathway related to amino acid/nitrogen metabolism or transport could be identified from the RNAseq data at 7HPI, which showed early responses to stress and regulation of transport processes. Alternatively, this time point might already be too late to capture any amino acid transport responses at the transcriptome level.

### The abscisic acid pathway is directly affected by over-expression of GDU1

The second notable effect of *GDU1* induction is the activation of ABA-related signaling and responses, occurring as soon as 12-24 HPI (Figures 3, Supplemental Figure 16). Interestingly, LOG2 was identified as a positive regulator of ABA signaling using a forward genetic screen (Kim and Kim, 2013). Our primary root growth assay under drought-mimic conditions also supports a positive role in ABA signaling not only for LOG2, but also for GDU1 (Figure 4B). While GDU1 over-expressors behave similarly to the wild type in this assay, they are hypersensitive to ABA in a germination test (Figure 4A), showing that both GDU1 and LOG2 are positive regulators of ABA/drought responses, potentially affecting overlapping processes. The working model for the GDU1-LOG2 complex can be expanded to a role in either sensitization of the plant to ABA or promoting ABA response. This model is consistent with the finding that RD21, encoding a Cys-protease induced by ABA and pathogen attack, is a substrate of LOG2 (Kim and Kim, 2013). RD21 plays a role in promoting programmed cell death, and is a target of pathogen effectors in some plant pathosystems (Lampl et al., 2013; Pogorelko et al., 2019). The combination of ABA sensitization and the stress induced by the modification in amino acid homeostasis could also trigger the ABA-mediated stress responses. The fact that the activity of the ABA pathway is transient upon *GDU1* induction, disappearing after ∼48 HPI (Figure 3) and not prevalently observed in *gdu1-1D*, could suggest an antagonistic effect of the ABA and SA pathways (see below) or the existence of a feedback loop on the signaling exerted by the GDU1-LOG2 complex.

The increase in amino acid content in leaves is concomitant with the activation of ABA signaling, occurring at about 24 HPI (Figure 1B). Treatment of plants with ABA leads to an increase in free amino acid content, originating from protein degradation rather than from *de novo* synthesis (Huang and Jander, 2017). The decreased expression of the genes involved in amino acid biosynthetic pathways at 25 HPI (Supplemental Figure 15A) is compatible with the hypothesis that the amino acids accumulating after *GDU1* induction also come from an ABA-initiated protein degradation. The over-accumulation of Pro, Leu, Val and Ile in *gdu1-1D* (Pilot et al., 2004) and at 96 HPI is in good agreement with an effect of ABA-mediated signaling, which leads to increased accumulation of these amino acids during salt and drought stresses (Urano et al., 2009; Kovacs et al., 2011; Huang and Jander, 2017). Alternatively, these increases in amino acid levels could originate from the enhancement of amino acid export rather than ABA signaling, supported by the drastic increase in amino acid concentration in the apoplasm after induction (Figure 1C and Supplemental Table 2). These responses could create a scarcity of amino acids in the cytoplasm, followed by modification of amino acid distribution in the cell or in the leaf. In response, protein degradation would be increased to replenish the stock of amino acids in the cytosol, leading to a global increase in amino acid content in the leaf without *de novo* synthesis. Rather than mutually exclusive, these scenarios could occur concomitantly with additive or synergistic effects.

Accumulation of the oxylipin hormone JA occurred concomitantly with ABA following induction of *GDU1* (Figure 3). JA has not been previously associated with amino acid homeostasis, and this potential connection is an interesting area for future exploration. JA biosynthesis and signaling declined as SA biosynthesis and signaling were activated, correlating with the well-documented antagonism between SA and JA signaling. However, it is intriguing to consider that accumulation of JA is an early event in SA-dependent systemic acquired resistance in Arabidopsis and JA has been proposed as a phloem-mobile systemic signal for SAR, in addition to Pip (Truman et al., 2007). Considering that activation of *GDU1* induces an SAR-like response, the DEX lines could provide a useful genetic tool for precise understanding of context-dependent interactions between JA, ABA, Pip and SA signaling in SAR.

### Disturbance in amino acid homeostasis triggers plant defense responses

Upon induction, SA and Pip accumulated and the corresponding pathways were activated after the ABA responses (Figure 3 and Supplemental Figure 3). Exogenous application of several amino acids could induce the expression of *PR1*, of which Cys, Gln, Pro, Gly, Phe and Ser were the most potent (Supplemental Figure 13). This result is in good agreement with previous studies, which showed that treating plants with Glu, Gln or Leu triggers defense responses (Hannah et al., 2010; Kan et al., 2015; Kadotani et al., 2016; Kan et al., 2017; Goto et al., 2020) and that amino acid disturbance of the knockout of the amino acid transporter AtLHT1 modulated SA responses (Liu et al., 2010). Increase in Lys content was larger and faster than any other amino acid upon *GDU1* induction, paralleled with increases in mRNA levels of Lys catabolic enzymes LKR-SDH and ALD1. *ALD1* encodes the enzyme catalyzing the first step of Pip synthesis (Navarova et al., 2012), a compound shown to orchestrate systemic acquired resistance and defense responses in concert with SA and ROS (Bernsdorff et al., 2016; Chen et al., 2018; Hartmann et al., 2018; Wang et al., 2018; Hartmann and Zeier, 2019). Our experiments with mutants deficient in either SA or Pip pathways indicate that two parallel processes could induce defense responses, one directly initiated by an amino acid-SA branch, and the other mediated by a Lys-Pip branch. The fact that gdu1-1D lines in which SA biosynthesis was abolished still over accumulated Pip and developed lesions (Figure 6) suggests that lesions in gdu1-1D are developed by Pip-mediated, SA-independent pathways. Suppression of most defense responses by expression of NahG (Heck et al., 2003) completely abolished lesion development (Figure 6A) and Pip (but not Lys) accumulation, suggesting that NahG activity inhibits the conversion from Lys to Pip through ALD1. In addition, induction of *GDU1* in the *ald1* background failed to accumulate ROS (Figure 6F), indicating a pivotal role of ALD1 and Pip in the development of lesions and defense responses in the Gdu1D phenotype. The induction of *PR1* expression by treatment of the *ald1* mutant with exogenous amino acids (Figure 6G) suggests that SA-dependent pathways are nevertheless triggered upon amino acid disturbance. No overlap between the ABA and SA responses was observed (Figure 3), prompting the hypothesis that ABA signaling is inhibited by the increase in the activity of the SA-mediated defense responses. Such antagonism is also evident when plants are simultaneously treated by biotic and abiotic stresses (Gupta et al., 2017), or in studying ABA receptors (Manohar et al., 2017).

### Model for the development of Gdu1D phenotype

One of the main advantages of chemically inducible systems (Moore et al., 2006) is to allow tightly regulated temporal and spatial misexpression of the gene of interest, which has been used to study plant development (Malinowski et al., 2011; Jiang and Berger, 2017; Tao et al., 2017; Balanza et al., 2018) and hormonal responses (Skalak et al., 2019), or to identify direct targets of transcription factors (Bargmann et al., 2013; Yamaguchi et al., 2015; Brooks et al., 2019). In the present study, we utilized the two-component pOp/LhGR system (Craft et al., 2005) to tackle a different problem common in plant biology, namely a pleiotropic phenotype. The chemically inducible gene expression system proved that the very first effect of *GDU1* over-expression is increased amino acid export, to a level of temporal precision that we never accomplished by comparing the wild type and constitutive over-expressors. This system allowed us to separate the primary and secondary effects and to formulate testable hypotheses on the causal relationships between them, providing a blueprint for understanding the role of unknown proteins, whole over-expression would lead to a recordable phenotype.

Our work leads to the model in which *GDU1* over-expression triggers in the following order: (1) Enhancement of amino acid export by controlling the activity of amino acid exporter(s). (2-1) Increase in amino acid export leading to more amino acids in the apoplasm, phloem and xylem, and disturbance in amino acid homeostasis. (2-2) Increase in amino acid levels, particularly in Lys, which is converted to Pip by ALD1. Stimulation of ABA signaling and responses mediated by the induction of LOG2 via interaction with GDU1. (3-1) As hypothesized earlier (Sonawala et al., 2018), increases in levels of apoplasmic amino acid could be a signature for the presence of a pathogen and trigger immune responses. (3-2) Defense responses involving Pip, SA and JA; Pip exacerbates *PR1* and other defense-related gene expression in both SA-dependent and SA-independent manners, and directly or indirectly mediates ROS accumulation. (4) The accumulation of SA in turn inhibits the activity of ABA and JA responses (Figure 7). The small size of the plants and Gln secretion in *gdu1-1D* could not be suppressed by inhibiting defense responses (Supplemental Figure 11A), but only by downregulating *LOG2* expression (Pratelli et al., 2012). Interestingly, loss of LOG2 activity does not completely suppress the Gdu1D phenotype (Supplemental Figures 6 and 7) suggesting that the effects of *GDU1* over-expression are mediated by LOG2 and other proteins, potentially the LOG2 homologs LULs (Pratelli et al., 2012). We hypothesize that disturbance in amino acid / nitrogen homeostasis is the reason for the growth reduction. The RNAseq analysis did not provide any clue that would help testing this hypothesis, possibly because these processes are masked by the large reprogramming of the transcriptome in response to stress.

Over-expression of *GDU1* leads to an interesting paradox: on the one hand, the increased amino acid content in leaves and various tissues provides an ideal source of carbon and nitrogen for pathogens, which could make plants more susceptible to pathogens (Zeier, 2013; Fagard et al., 2014; Mur et al., 2017; Sun et al., 2020). On the other hand, the augmented SA- and Pip-mediated defense response increases plant disease resistance. More research will be needed to tell which process will prevail (if any), based on assessment of disease susceptibility of *gdu1-1D* compared to the wild type, taking into account that the effect could be different for distinct pathogens, at different time points following induction.

## Conclusions

Our work provides solid evidence that over-expression of *GDU1* triggers two parallel pathways most likely involving the LOG2 ubiquitin ligase: post-translational regulation of amino acid export, and ABA- (and possibly JA-) mediated stress responses first visible at the level of the transcriptome. This brings further evidence for an interesting relationship between the co-regulation of ABA signaling and nitrogen metabolism, in which the GDU1-LOG2 complex could play a critical role. GDU1-mediated disturbance in amino acid homeostasis, independent on pathogen attack, triggers plant immune responses involving Pip, SA, and JA. *GDU1* over-expressors and the induction system would provide a unique resource to study the interaction between amino acid homeostasis, stress-related phytohormones and plant immunity.

## MATERIAL AND METHODS

### Plant material and growth

*Arabidopsis thaliana* plants were grown under 120 μmol.m^-2^.s^-1^, 22°C, 16/8 h light/dark cycle on soil (Mix of Sunshine Mix 1 and Pro-mix HP at a 1:1 ratio) and were watered from below with 300 mg/l Miracle-Gro Fertilizer (24/8/16 NPK; Scotts, Marysville, OH, USA). g*du1-3* (SALK_132115) and *ald1* (SALK_007673; (Alonso et al., 2003)) were obtained from the Arabidopsis Biological Resource Center. *glr3*.*3-1* and *glr3*.*4-1* were gifts from Dr. Edgar Spalding (University of Wisconsin at Madison, USA). pPR1-GUS was a gift from Dr. John McDowell (Virginia Tech, USA). The pOp/LhGR plasmid and the control line 4c-S7 were obtained from Dr. Ian Moore (Oxford, UK). *Arabidopsis thaliana* plants were transformed by the floral dip method (Clough and Bent, 1998) using *Agrobacterium tumefaciens* GV3101 (pMP90). Expression of *GDU1* in soil-grown plants was induced by spraying with a solution composed of 100 μM dexamethasone (100 mM stock solution in DMSO) and 300 ppm silwet-77. For phytohormones, RNA, and metabolite analyses, each biological sample corresponded to 3-4 adult leaves from a single plant. For other experiments, plants were grown on solid half-strength Murashige and Skoog (MS) medium supplemented with 1% sucrose for 7 days, about 6-8 seedlings were transferred to one well of a 12-well plate containing 3 ml of ½ MS + 1% sucrose grown for four more days in a growth chamber (same conditions as above) with gentle shaking (40 rpm). Induction was performed with 30 μM dexamethasone and a biological replicate corresponded to plants from one well. pPR1-GUS seeds were grown on solid ½ MS + 1% sucrose medium 8 days, six seedlings were transferred into a well of a 24-well plate filled with 2 ml of liquid ½ MS + 1% sucrose and grown for three additional days under gentle shaking. Amino acids and other compounds were added at the indicated concentration to trigger *GUS* expression from stock solutions, staining was performed 2 days after treatment. The negative control was treated with 0.025% DMSO final (highest volume DMSO). For germination assays, about 150 seeds for each genotype were sowed on solid ½ MS + 1% sucrose medium supplemented with ABA at the indicated concentrations. After three-day stratification at 4°C, dishes were transferred to a growth chamber under 120 μmol.m^-2^.s^- 1^, 22°C, 16/8 h light/dark cycle, and seed germination rate (cotyledon greening) was counted five days later. For root elongation assays on PEG, seeds were grown on solid ½ MS, 1% sucrose medium for six days. Seedlings were transferred to solid ½ MS + 1% sucrose medium ± polyethylene glycol PEG 8000, as described (van der Weele, 2000). Briefly, 250 g of PEG 8000 was dissolved in 500 ml of the liquid medium (½ MS + 1% sucrose), and filtered through 0.22 μm PES filter. Roughly 30 ml of PEG solution was poured on top of an equal volume of the solid medium. After 24 h, the solution was discarded, and the plate was used for the experiment. The average water potential of media with and without PEG 8000 was −0.88 MPa and −0.04 MPa, respectively (measured using a Decagon WP4 dew point potentiometer). Seedlings were then grown vertically under 120 μmol.m^-2^.s^-1^, 22°C, 16/8 h light/dark cycle for a week; dishes were scanned, and primary root length was measured using ImageJ (Schneider et al., 2012). Mutant lines used in this study are listed in Supplemental Table 5. Samples collected from the same plants and at the same time for different analyses and assays are indicated in figure legends.

### Amino acid uptake in seedlings

Measurements of amino acid transport were performed as previously described (Pratelli et al., 2010), with the following modifications for the data presented in Supplemental Figures 7 and 10: plants were grown for 7 days on solid, half-strength MS medium containing 1% sucrose, and transferred to 1 ml of MS medium containing 1% sucrose for five more days in a 24-well plate without shaking; the solution was replaced by fresh medium, containing unlabeled Gln and 0.5 µl of labeled ^3^H-Gln (18.5 kBq total).

### Nucleic acid manipulation and RNA seq analysis

Details are given in Supplemental Text 3.

### Metabolite and hormone level measurements

For amino acid analyses, samples were lyophilized, and homogenized with two 3 mm glass beads in a bead beater twice for 60 s at 60Hz (Mini-Beadbeater-96, Biospec, USA). About 1.5 mg homogenized samples were transferred to a tube containing 10 μl of 2 mM norvaline previously dried as an internal standard. Samples were extracted twice in 200 μl 10 mM HCl and 200 µl chloroform. The supernatants were pooled and transferred to a fresh tube for UPLC analysis. Derivatization and analysis were performed as described (Collakova et al., 2013). Hormone analyses in leaves were performed essentially as described (Forcat et al., 2008), with the following modifications. Samples were lyophilized, and homogenized with two 3 mm glass beads, by shaking in a bead beater twice for 60 sec at 60Hz. About 10 mg of homogenized samples were transferred to a tube and extracted twice in 400 μl of a solution composed of 10% methanol and 1% acetic acid in water. The supernatants were pooled and analyzed by LC/MS-MS (see Supplemental Text 3).

For apoplastic washing fluid collection, *Arabidopsis* plants were grown on soil for four weeks. 6-8 adult leaves were collected from each plant by cutting from the base of the petiole, and infiltrated with a solution containing 240 mM sorbitol and 6 mg.L^-1^ Lucifer Yellow CH dipotassium salt (LYCH, Sigma), which is used as a tracer to normalize apoplastic fluids (Derrick et al., 1992). The leaves were then stacked and rolled into a 5 ml tip, inserted into a 15 ml conical tube. The apoplastic wash was recovered by centrifugation at 22°C for 5 min at 400xg. LYCH concentration was assessed in a microplate reader (Synergy4, BioTek, USA) using an excitation of 428 nm and emission of 536 nm at room temperature. The intactness of the apoplastic wash fluids was confirmed by measuring hexose phosphate isomerase activity as described (Dannel et al., 1995).

### Histochemical staining

GUS activity was revealed by histochemical staining, performed as described (Lagarde et al., 1996). ROS staining by diaminobenzidine and cell death staining were performed as described (McDowell et al., 2011).

### Reactive oxygen species measurement

Measurement of ROS was performed as described (Umbach et al., 2012) with the following modifications. Leaves were excised and incubated in 3 ml of a solution containing 20 μM 2’,7’-dichlorofluorescin diacetate, 10% MS and 0.1% Tween-20 in a 12-well plate in the dark for 30 min at room temperature. Leaves were transferred to a fresh tube, dried at 80°C and weighed. The liquid medium was separated in four 200 µl aliquots, transferred to a 96-well plate and the fluorescence was measured with excitation at 488 nm and emission at 525 nm using a Synergy4 microplate micro plate reader.

## Supplemental Material

Supplemental Figure 1. Root length analysis of Ile tolerance of the DEX lines.

Supplemental Figure 2. Phenotype of the *gdu1-1D* and Col-7 leaves.

Supplemental Figure 3. Expression analysis by qRT-PCR of marker genes at different time points following GDU1 induction.

Supplemental Figure 4. Gln uptake and efflux of plants treated by ABA, SA or Pip.

Supplemental Figure 5. Response of the gdu1-3 knockout mutant to ABA.

Supplemental Figure 6. Amino acid content in leaves of plants with modified expression in GDU1 and LOG2.

Supplemental Figure 7: Gln uptake and efflux in plants with misexpression in GDU1 and LOG2.

Supplemental Figure 8. Levels of GDU1 and PR1 transcripts in gdu1-1D plants, in which defense responses have been suppressed by genetic approaches.

Supplemental Figure 9. Phytohormone content in gdu1-1D plants in which defense responses have been suppressed by genetic approaches.

Supplemental Figure 10. Gln uptake of gdu1-1D plants in which defense responses have been suppressed by genetic approaches.

Supplemental Figure 11. Picture of gdu1-1D plants in which defense responses have been suppressed by genetic approaches.

Supplemental Figure 12. PR1 and GDU1 expression in mutants harboring a dexamethasone-inducible expression of GDU1 construct after treatments with dexamethasone.

Supplemental Figure 13. GUS activity in pPR1-GUS plants treated with various amino acids, and compounds.

Suppl. figure 14. Analysis of transcriptomic changes for induced DEX plants and the gdu1-1D mutant.

Supplemental Figure 15. Mapman analysis of transcriptome response to GDU1 over-expression.

Supplemental Figure 16. Average fold changes of transcript levels of 40 genes, used as a marker for each indicated treatment.

Supplemental Figure 17. Analysis of effects of suppressing the expression of GLRs on the response to Ser and to GDU1 induction.

Supplemental Figure 18. Effect of treatments with GLR antagonists on dexamethasone-treated 4c-S7 and DEX1.

Supplemental Figure 19. Maps of constructs used in this study.

Supplemental File 1. RNAseq output and clustering analysis.

Supplemental File 2. GO analysis of each gene clusters.

Supplemental File 3: Analysis of the clusters using the signature tool from Genevestigator

Supplemental File 4: Marker search using Genevestigator for various stresses and pathways.

Supplemental File 5: Fold changes of stress and nitrogen metabolism genes in response to GDU1 induction.

Supplemental File 6: Signature analysis of the nitrogen metabolism genes using Genevestigator.

Supplemental Table 1: Amino acid content in leaves of induced and non-induced plants.

Supplemental Table 2: Amino acid composition in the apoplasm wash fluid.

Supplemental Table 3: Marker genes used for qRT-PCR analysis.

Supplemental Table 4: Amino acid content in leaves of SA-related crosses and transformations.

Supplemental Table 5: Plant lines used in this study.

Supplemental Table 6: Sequences of the oligonucleotides used for this work. Supplemental Table 7: TOPHAT statistics.

Supplemental Text 1: Untargeted transcriptomic analysis unravels the transition from ABA to defense responses.

Supplemental Text 2: GLRs are not positive regulators of the events downstream from GDU1 induction.

Supplemental Text 3: Supplemental Material and Methods

## ACCESSION NUMBERS

Sequence data from this article can be found in the EMBL/GenBank data libraries under accession numbers AT4G31730 (GDU1), AT3G09770 (LOG2). RNAseq output data can be found in the GEO data library under the accession number XXXXXX.

## ACKNOWLEDGEMENTS

The authors thank Dr. Ryan Stewart (Virginia Tech, USA) and Dr. Josh Heitman (North Carolina State University, USA) for water potential readings, Dr. Edgar Spalding (University of Wisconsin at Madison, USA) for the *glr3*.*3-1, glr3*.*4-1* lines, Dr. John McDowell (Virginia Tech, USA) for the pPR1-GUS line, and Dr. Ian Moore (University of Oxford, UK) for sharing the dexamethasone inducible constructs and plant lines, Dr. Cynthia Denbow and Dr. John McDowell for critical reading of the manuscript.

